# Spaces and sequences in the hippocampus: a homological perspective

**DOI:** 10.1101/2025.02.08.637255

**Authors:** A. Babichev, V. Vashin, Y. Dabaghian

## Abstract

Topological techniques have become a popular tool for studying information flows in neural networks. In particular, simplicial homology theory is used to analyze how cognitive representations of space emerge from large conglomerates of independent neuronal contributions. Meanwhile, a growing number of studies suggest that many cognitive functions are sustained by serial patterns of activity. Here, we investigate stashes of such patterns using *path homology theory*—an impartial, universal approach that does not require *a priori* assumptions about the sequences’ nature, functionality, underlying mechanisms, or other contexts.

We focus on the hippocampus—a key enabler of learning and memory in mammalian brains—and quantify the ordinal arrangement of its activity similarly to how its topology has previously been studied in terms of simplicial homologies. The results reveal that the vast majority of sequences produced during spatial navigation are structurally equivalent to one another. Only a few classes of distinct sequences form an ordinal schema of serial activity that remains stable as the pool of sequences consolidates. Importantly, the structure of both maps is upheld by combinations of short sequences, suggesting that brief activity motifs dominate physiological computations.

This ordinal organization emerges and stabilizes on timescales characteristic of spatial learning, displaying similar dynamics. Yet, the ordinal maps generally do not reflect topological affinities—spatial and sequential analyses address qualitatively different aspects of spike flows, representing two complementary formats of information processing.

## I. INTRODUCTION

One of the central tasks in neuroscience is to explain learning and memory through neuronal activity [1]. A key role in enabling these phenomena is played by the hippocampus—one of the brain’s fundamental regions, both functionally and phylogenetically. Broadly, there are currently two perspectives on the organization of hippocampal neuronal activity. On the one hand, it is believed that memory episodes may be encoded and invoked by the firings of individual cells and their combinations—*cell assemblies*^1^. For example, hippocampal neurons, called place cells, learn to recognize specific areas within an environment—their respective place fields. Subsequent firing of these cells can be triggered not only by the animal’s physical visits to these fields but also by internally generated activity: recalls of previous visits or anticipations of upcoming ones [2–5]. Suppressing particular cells’ activity using electrophysiological or optogenetic tools can prevent the animal from venturing into the corresponding fields, whereas stimulation can prompt such visits [6, 7]. Curiously, hippocampal cells may also learn to selectively respond to specific odors, visual cues, objects, memory episodes, and more, thus linking spatial and mnemonic information [8–12].

On the other hand, the hippocampus also enables animals to understand event sequences, plan series of actions, learn cue arrays, and more [13–15]. For instance, rats with hippocampal lesions can recognize odors but struggle to memorize their order [13]; lesioned monkeys have difficulty learning new stimulus sequences [13, 16]; humans with hippocampal damage lose the ability to rank the positions of objects in learned sequences [17–20], *etc*. Physiologically, sequential learning and recall are supported by consecutively firing cells and their assemblies; accordingly, the temporal order of serial spiking activity currently receives significant attention [21–25].

Altogether, the hippocampus appears to provide a scaffold for multimodal relationships among memory elements and to enable sequential activity as a dynamic expression of that scaffold. In particular, a stable configuration of the hippocampal network may reflect the spatial structure of the environment and serve as a template for representing sequences of events, of which spatial trajectories are a salient, experimentally accessible example [25, 26]. Importantly, this template can accommodate a wide range of details through interactions with other brain regions. For example, memorized sequences can scale to different physical sizes and temporal durations, adapting to specific tasks and conditions [27–29]. Moreover, this scaling can be modulated by cognitive and behavioral context; items may be subjectively judged as ‘closer’ or ‘farther apart’ depending on circumstances [30]. Similarly, hippocampal cells preserve their relative spiking order amid gradual geometric transformations of the navigated environment [31–37], further pointing to the qualitative nature of the encoded map [37–40].

These properties suggest that the hippocampal network encodes a qualitative, relational framework. Prior models of this framework represented patterns of simultaneous coactivity using topological constructions—simplicial complexes, and analyzed their boundaries, voids, *etc*., with various forms of simplicial homology, while not addressing the ordinal aspects of firing [40–51]. To address these, we employ path homology theory, which translates the network-enforced causal grammar of spiking into an ordinal structure: how admissible sequences interact and compose, the “navigability” of directed routes across scales, *etc*. [52–56].

The two homology theories have different purviews, *e.g*., path homology ignores all-to-all coactivity cliques and vanishes on acyclic directed graphs, whereas simplicial homology can detect “holes” in the corresponding complexes but is insensitive to the existence of long navigable sequences or to their compositions beyond simplex boundaries. Considering both perspectives puts complementary aspects of information processing in the hippocampal network on equal footing, thereby qualitatively broadening the topological framework [37, 38, 40, 80].

The paper is organized as follows. Section II discusses current topological models of hippocampal activity and outlines path homology theory along with its connections to physiology. The results of path-homological analyses of simulated neuronal dynamics are presented in Section III and discussed in Section IV.

## II. TOPOLOGICAL METHODS

### A. Simplicial schema of cognitive map

A cell assembly is a transient group of neurons that work together^2^ to elicit responses from downstream “readout” neurons [24]. A given neuron, *c*_*i*_, may participate in multiple assemblies, meaning the cell assemblies are interconnected. Formally, an assembly can be viewed as an abstract simplex (Fig.1A),

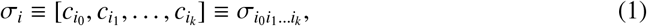

with the subsimplexes corresponding to independently activating “subassemblies” of *σ*_*i*_ [58–60]. The combinations of cells activated by time *t* form a simplicial coactivity complex,

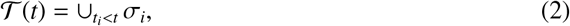

which provides a collective representation of the information encoded by individual spiking units [48]. The shape of the coactivity complex reflects the overall, large-scale structure of information and exhibits properties manifesting at the cognitive level. For instance, such complexes capture the shape of the navigated environment: temporally filtered persistent simplicial homologies of 𝒯(*t*) evolve to match the homological structure of the environment, *H*_∗_(𝒯(*t*)) = *H*_∗_(ℰ), *t* > *T*_min_ (for definitions, see Sec.VI). The minimum time required for this process, *T*_min_(𝒯), approximates the physiological *learning time* [40, 42–49].

Such a simplicial schema of neuronal activity aids in interpreting neurophysiological computations. For example, the assemblies igniting along a navigated path, *γ*, induce a series of consecutively activating simplexes that form a representation of *γ*—a simplicial trajectory,

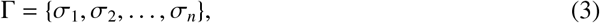

embedded in 𝒯. Such trajectories not only represent ongoing behavior [61–63], but they also enable the reconstruction of an animal’s past navigational experiences and the prediction of upcoming, planned journeys [13]. Even the existence of place fields can be viewed as an intrinsic, topological property of the coactivity complex [64, 65].

The structure of simplicial complexes is derived from the adjacencies of their simplexes. Algorithmically, this is achieved by introducing an operator, *∂*, which splits the boundary of each simplex (1) into facets, as illustrated in Fig. 1B,

**FIG. 1.**
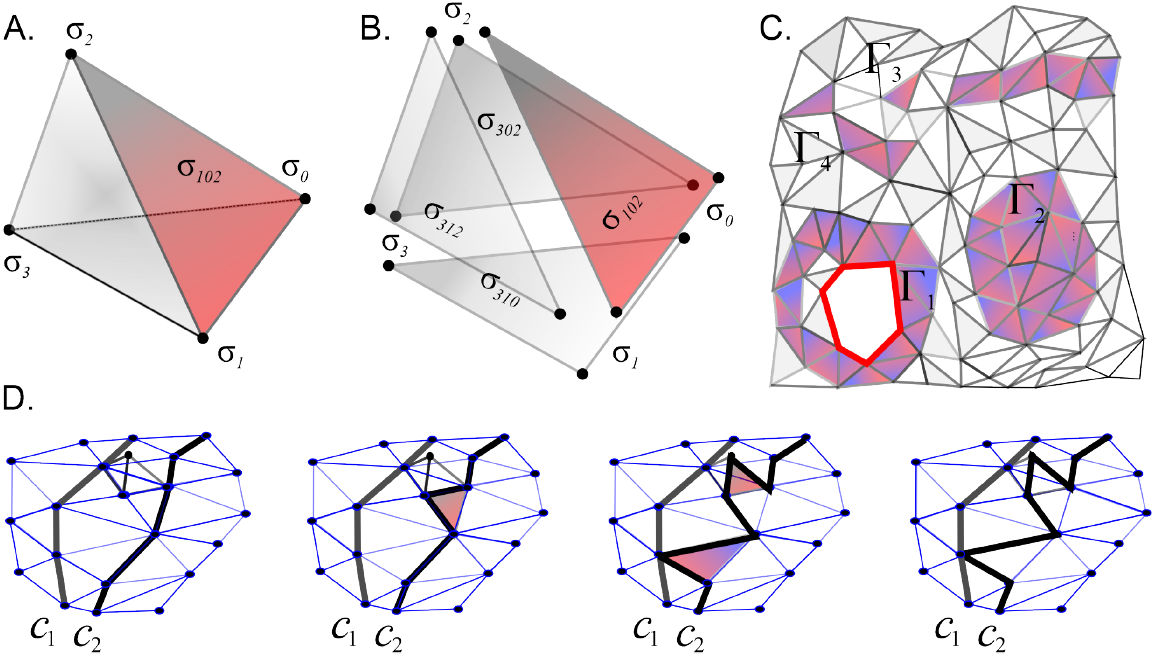
Simplexes and simplicial chains. **A**. Simplexes are *k*-dimensional polytopes spanned by *k* + 1 vertices. Shown is a three-dimensional (3*D*) simplex—a tetrahedron. **B**. The boundary of a 3*D* simplex is a combination of its two-dimensional (2*D*) facets. **C**. Simplexes with matching boundaries form a simplicial complex. A chain in such a complex is an arbitrary combination of simplexes, counted with coefficients from a field 𝕂. Here colored simplexes illustrate 2*D* chains connected (Γ_1_, Γ_2_, Γ_3_) and disconnected (Γ_4_), contractible (Γ_2_, Γ_4_) and topologically nontrivial—a 1*D* cycle (*c*_1_, red). **D**. Two homologous one-dimensional (1*D*) chains, *c*_1_ and *c*_2_, highlighted by thick black lines. The panels show a series of discrete deformations that transform one chain into the other, produced by “snapping” segments of *c*_2_ over the boundaries of adjacent (red) simplexes.

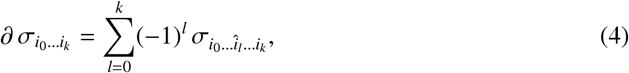

where the hat over an index *i*_*l*_ denotes the omitted vertex 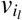 [58–60]. The key objects in the theory are simplex chains—collections of simplexes that span the complex and collectively capture its shape. For instance, simplicial trajectories (3) can be viewed as contiguous chains of abutting simplexes.

If a simplicial chain borders a simplex of higher dimensionality, its segments can be “snapped over” the boundary of that simplex, producing a chain deformation (Fig.1D). Correspondingly, if two chains, *s*_1_ and *s*_2_, differ by a collection of full boundaries— they may be considered equivalent, or homologous (Sec.VI). Sets of homologous cycles correspond to elements of the complex’s topological structure, such as holes, cavities, pieces, tunnels, and so forth [58–60]. Specifically, classes of equivalent simplicial trajectories (3) in representable coactivity complexes mirror the structure of the environment covered by the place fields, including obstacles encircled by the rat or shortcuts the rat can take [40–44].

Algebraically, sets of homologous chains form abelian groups or vector spaces, depending on the coefficients used to count the simplexes. With coefficients from an algebraic field, 𝕂, one obtains *n*+1 vector spaces, one for each dimension: *H*_0_(𝒯, 𝕂), *H*_1_(𝒯, 𝕂), …, *H*_*n*_(𝒯, 𝕂), commonly referred to as the simplicial homologies of 𝒯^3^. The dimensionalities of these homology spaces, *b*_*k*_ = dim(*H*_*k*_(𝒯, 𝕂)), known as Betti numbers, count connectivity components (*β*_0_), holes and tunnels (*β*_1_), cavities (*β*_2_), and so on.

In practice, the hippocampal coactivity complex is often constructed as the clique complex associated with the graph of coactivity, 𝒢, also known as the cognitive graph [66–72]. The nodes of this graph correspond to principal cells, and its links represent either physiological or functional connections between them. If these connections are directed (*e.g*., extending from presynaptic to postsynaptic cells), the graph is directed and referred to as a *digraph*. Otherwise, if the connections represent undirected coupling, *e.g*., rate of co-firing, the coactivity graph is undirected.

In applications, specific constructions of cliques may be used to capture different physiological phenomena. For instance, cliques formed over time from frequently co-occurring lower-order coactivities (*e.g*., pairs or triples of cells) can model input-integrating cell assemblies, whereas large groups of simultaneously co-firing cells can represent coincidence-detecting cell assemblies. Both mechanisms have been experimentally identified, making the corresponding coactivity complexes viable functional representations of physiological coactivity pattern [42–49].

### B. Sequential schema of neuronal activity

If ordered sequences of neurons’ firings, rather than nearly-independently igniting cell groups, serve as functional units of neuronal activity, then the approach described above requires significant modifications. Let us represent the population of active units (cells or their assemblies) by a set of vertices, *V* = {*v*_1_, *v*_2_, …, *v*_*n*_}, and consider sequences of various lengths,

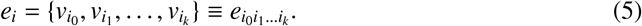

Following the terminology of path homology theory, we will also refer to (5) as an *elementary path* running through the set *V* [52, 53]. A simplicial path (3) is one example of an elementary path comprised of igniting cell assemblies, while a sequence of individual neurons activated autonomously in the animal’s brain *(e.g*., during sleep) is another [73–75]. Note that, since the firing units in (5) spike in order at times 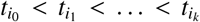, each sequence (5) is characterized by a specific start and completion time. The set of spiking sequences observed up to time *t* defines a *path complex*

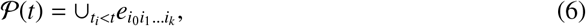

which is the path-analogue of 𝒯(*t*) (Fig. 2A), described by an alternative homology theory [52– 55]. The only requirement for the collection of paths (6) is that if a path *e*_*i*_ is included, then the paths obtained by “plucking” the ends of *e*_*i*_ must also belong to 𝒫(*t*) (Fig. 2B). In other words, given a cell sequence (5), its shorter contiguous subsequences are assumed to contribute to the informational framework encoded by 𝒫(*t*)—a physiologically natural assumption.

**FIG. 2.**
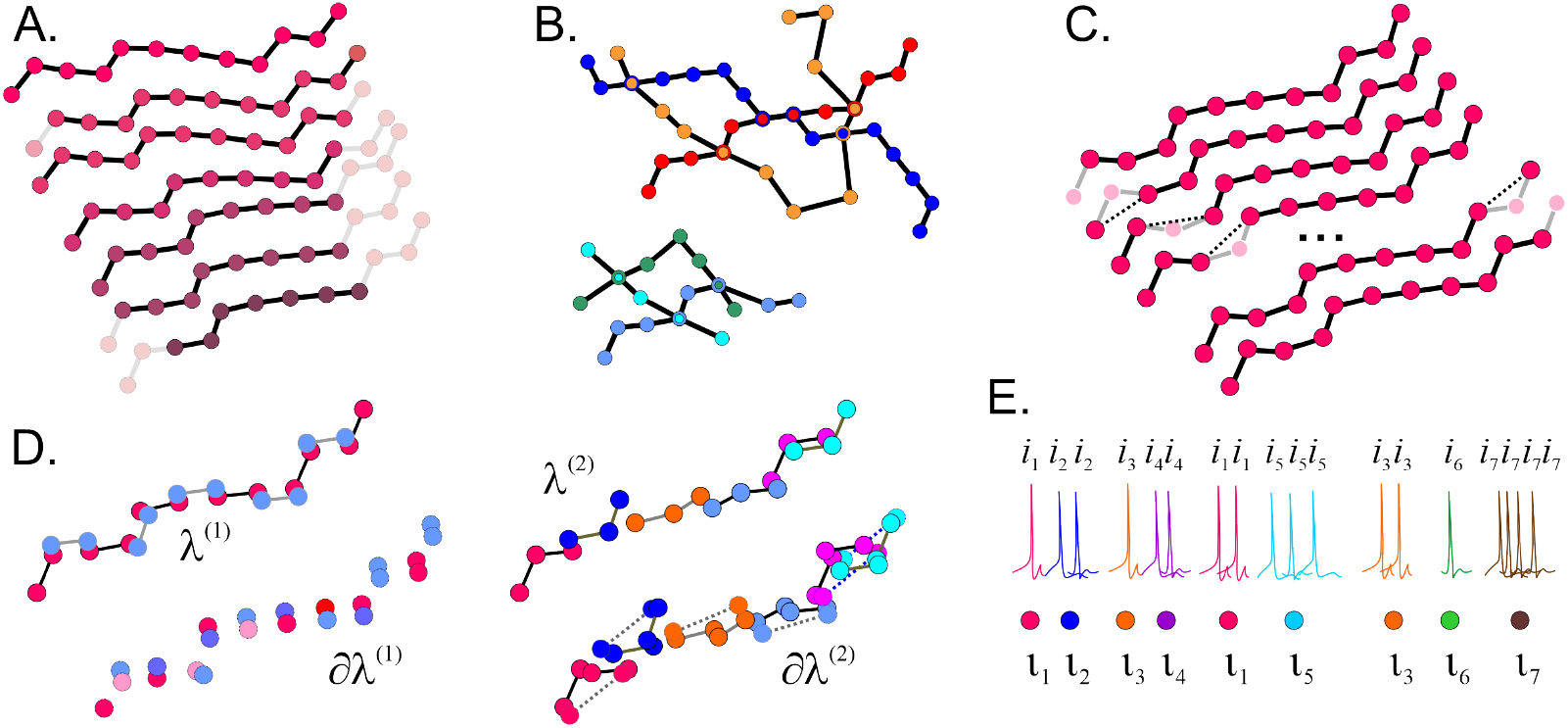
Path, path chains, path complexes. **A**. An elementary path *e* of length 12 and examples of shorter paths obtained by “plucking” its end vertices. All such truncated paths must be included in any path complex that contains *e*. **B**. A path complex with two components and six maximal *simple* elementary paths (without vertex repetitions). The presence of all subpaths obtained by plucking these longest paths is implied. **C**. The path-boundary (7) of the sequence *e* illustrated on the top of panel A. Note that only the first and the last paths shown on this panel also appear on panel A. **D**. A sequence of *k* elements can be viewed as a combination of pairwise links (a length-one path chain, λ^(1)^, left panel) or triples of consecutive vertices (a length-two path chain, λ^(2)^, right panel), *etc*. The boundaries of these chains, *∂* λ^(1)^ and *∂* λ^(2)^, are shown below. **E**. A collection of bursting neurons *ι*_1_, *ι*_2_, …, *ι*_7_ produces sequences of spikes marked by repeated indices *i*_*k*_: neuron *ι*_2_ yields two spikes in a burst, neuron *ι*_5_ yields three spikes, and so on. Regularized sequence reveals the order in which the neurons fire (bottom).

Note that constructing a path complex, *e.g*., from a dataset of spiking time series (spike trains), requires that all elementary paths be specified explicitly: their existence cannot be presumed. In particular, one should not assume that paths connect all vertices (neurons) or follow any prescribed configuration or track: they may or may not be present. Nevertheless, for certain sequence grammars it may be possible to identify a generative template” [74–76]. For instance, if a path complex 𝒫 contains paths assembled from specified pairs of vertices, then one can induce 𝒫_*G*_ by traversals of a graph *G* in which those pairs are joined by edges [52, 53]. In such cases, one may treat a finite set of observed paths as partial and consider the “ultimate” path complex, consisting of all paths consistent with the graph’s connectivity—the entire pool of admissible vertex sequences. Physiologically, such a graph may arise, for example, as the synaptic wiring of the underlying network or as a coactivity graph [76]. In what follows, we focus on these *graph-representable* path complexes, though the theory permits more general constructions.

The homological description of path complexes develops similarly to simplicial homology theory. Viewing an elementary path (5) of length *k* as analogous to a *k*-dimensional simplex, one can construct its “structural boundary” as the formal alternating sum,

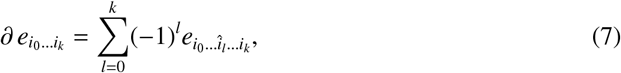

where 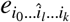 runs from 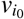 to 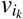, omitting the vertex 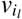 (Fig. 2C). Thus, the “boundary” of a path of length *k* is a combination of its (*k* + 1) subpaths, each of length *k* − 1.

The next step is to consider *path chains*—arbitrary combinations of elementary paths—and to view two chains as homologous to one another if their difference is the boundary of another path chain, in the sense of formula (7) [52, 53]. The resulting equivalence classes, properly counted, form vector spaces called the *path homologies* of a path complex 𝒫, which describe its structure just as simplicial homologies describe simplicial complexes [52–55].

This outline requires several stipulations. First, unlike the subsimplexes on the right-hand side of (4), which architecturally belong to 𝒯(*t*), the “path-facets” on the right-hand side of (7) may not fit the path complex, meaning that formula (7) may be ill-defined. For example, for a path complex 𝒫_*e*_ that contains a single linear path (5) and its required “truncations,” the formula (7) generates subpaths that skip intermediate vertices and therefore do not belong to 𝒫_*e*_ (Fig. 2C).

Path homology theory solves this problem by restricting the operations to the *allowed* paths— those contained in 𝒫, and, within these, to those whose boundaries are also allowed—the *operational* paths. The path homology groups are then defined as cycles modulo boundaries within the operational paths, denoted here as 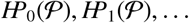.

For details, we refer the reader to the original publications [52–55] and to Sec. VI. As an illustrative example, consider a digraph with three vertices *v*_1_, *v*_2_, *v*_3_ and edges *v*_1_ →*v*_2_, *v*_2_ →*v*_3_, *v*_3_ →*v*_1_. The associated complex 𝒫_*G*_ consists of all admissible vertex sequences compatible with these edges: the 0-paths are the vertexes; the 1-paths are the three directed edges; the admissible paths are *v*_1_*v*_2_*v*_3_, *v*_2_*v*_3_*v*_1_, and *v*_3_*v*_1_*v*_2_; longer admissible paths are obtained by concatenation. In particular, shortcuts such as *v*_1_*v*_3_ (or *v*_2_*v*_1_, *v*_3_*v*_2_) are not admissible (Fig. 3A). All 0^th^ and 1^st^ order paths are allowed and operational—the differential (7) acts freely on *P*_0_ and *P*_1_, *e.g*., *∂*(*v*_1_*v*_2_) = *v*_2_ − *v*_1_.

**FIG. 3.**
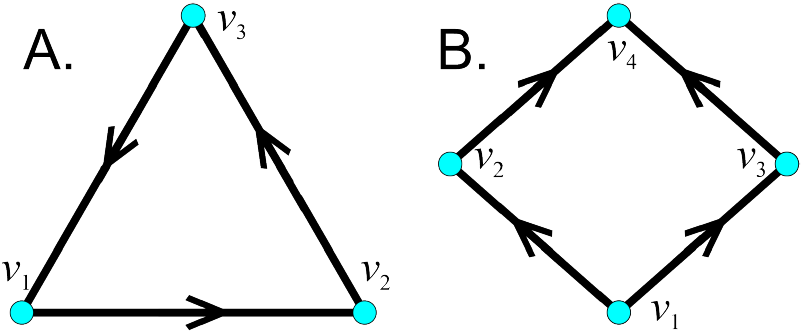
Comparison of simplicial and path homologies. **A**. Directed 3-cycle. Path homology detects a nontrivial 1-class: 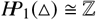 expressing the existence of a genuine directed loop—non-contractible flow. Undirected clique complex of the underlying graph is a filled triangle (a 2-simplex), hence *H*_1_(Δ) = ℤ, since, simplicially, the 1-cycle is “filled” and disappears. Directed clique complexes typically exclude a 2-simplex here (no acyclic ordering), so a 1-cycle may persist; nevertheless, the meaning remains co-incidence-based, not path-compositional. **B**. Acyclic digraph, with *v*_2_ and *v*_3_ unconnected. Its path homologies for *k* ≥ 1 vanish, 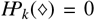, indicating no directed flow obstruction. Note that undirected clique complex of the underlying graph contains a 4-cycle *v*_1_−*v*_2_−*v*_4_−*v*_3_−*v*_1_, yielding a nontrivial simplicial homology *H*_1_(◊); which signals a “hole” that is irrelevant for directed reachability.

The 2^d^-order paths introduce complications, *e.g*., the differential

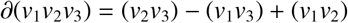

contains a component, *v*_1_*v*_3_, which is not a part of the complex, as there is no edge *v*_1_ → *v*_3_. Path homology theory prescribes simply to drop this term, and obtain

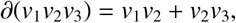

and, similarly, use *∂*_2_(*v*_2_*v*_3_*v*_1_) = *v*_2_*v*_3_ + *v*_3_*v*_1_ and *∂*_2_(*v*_3_*v*_1_*v*_2_) = *v*_3_*v*_1_ + *v*_1_*v*_2_. The resulting paths are not operational: using the rule (7), one finds that the boundary of the boundary chain *v*_1_*v*_2_ + *v*_2_*v*_3_ is (*v*_2_ − *v*_1_) + (*v*_3_ − *v*_2_) = *v*_3_ − *v*_1_ ≠ 0, and similar problems occur for chains built from *v*_2_*v*_3_*v*_1_ and *v*_3_*v*_1_*v*_2_. Thus there are no nontrivial second-order operational paths in this example. As a result, the first path homology group is simply the space of closed 1-chains. A direct check shows that the only nontrivial closed 1-chain (up to scaling) is

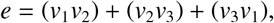

so 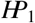 is one-dimensional and represented by this cycle.

The unusual properties of the path boundary illustrated above are important: if paths responded to the boundary operator (7) in the same way that simplexes respond to the boundary decomposition (4), the two homology theories would emulate one another. Instead, the principles for identifying structurally equivalent serial patterns are genuinely different from the principles establishing simplicial equivalence, pointing to qualitatively distinct organizations of spatial and ordinal frameworks.

Of course, paths that behave like simplexes do exist, but they are non-generic and consist only of those paths that traverse simplex vertices. A complex 𝒫 comprised exclusively of such “folded-into-asimplex” paths, which contain *all* boundary subpaths of *all* of its elementary paths, is structurally similar to a simplicial complex and is referred to as *perfect* in [52–55]. However, most path complexes are far from perfect; that is, boundary subpaths of most of their elementary paths are missing, resulting in path homologies that differ substantially from the simplicial homologies of the underlying graph’s clique complex.

The second point is that there are multiple ways to interpret a given firing series as a path. A string of *k* vertices can be viewed not only as a single path of length *k*, but also as a combination of 1-vertex paths—individual nodes (a 0-chain, *λ*^(0)^), a collection of 2-vertex paths (a 1-chain of graph links, *λ*^(1)^), or as sequences of 3-vertex paths, *i.e*., link pairs (a 2-chain, *λ*^(2)^), link triples (a 3-chain, *λ*^(3)^), and so on, with all shorter constituents overlapping in arbitrary ways (Fig. 2D). As it turns out, reasoning in terms of such “path chains,” rather than individual paths, allows for rectifying the unencumbered boundary operator (7) and transforms any collection of paths into a self-contained complex [52–55]. In the previous example, a balanced chain (II B) of all three 1paths has a vanishing boundary, and no illegal shortcuts. Thus, *e*, as a combination, is operational and serves a generator of homological loops.

Third, paths that repeatedly traverse the same vertices, such as 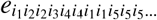, are possible but often redundant in practical analyses. For example, consider a complex where the elementary paths correspond to sequences of spikes, each indexed by the neuron that produced it. Since neurons tend to “burst,” *i.e*., fire spikes in rapid succession, such paths will contain multiple repeating indices (Fig.2E, [77, 78]). In analyses focusing on the order of neuron activation, these repetitions are uninformative and should be factored out. Conveniently, path homology theory allows for the exclusion of index repetitions by reducing each path to its unique *regular representative* with distinct consecutive vertices [52, 53]. At the graph level, this corresponds to removing trivial loops, *e*_*ii*_. The resulting paths form *regular path complexes*, described by *regular path homologies*. Below we consider only these regular path complexes and omit the term “regular” (see Sec.VI).

These and other properties point that simplicial homology—whether on undirected or directed complexes—and path homology approaches have distinct purviews. The former quantifies the topology of simplicial complexes, including clique complexes built on coactivity graphs, whereas the latter quantifies directed “navigability” and the composition of admissible paths (Fig. 3B). In neuronal terms, simplicial homology captures higher-order coactivity patterns: a clique (and its associated simplex) asserts that the units at its vertices can exhibit pairwise coactivity within an appropriate temporal window [42, 43]. A “void of coactivity” indicates a structured absence of joint activation, *e.g*., disconnected clusters that coactivate internally but not with each other induce *H*_0_; closed chains of pairwise coactivity that wrap around a loop without being “capped” by coactive triples produce *H*_1_; cavities bounded by many coactive triples but lacking quadruple coactivity to fill the interior generate *H*_2_, *etc*. In general, a nonzero *H*_*k*_(𝒯) certifies that some (*k* + 1)-order coactivity pattern is globally obstructed despite abundant lower-order coactivity. However, these homologies are largely insensitive to the existence of navigable sequences or to the algebra of their compositions. By contrast, path homology groups 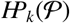 detect obstructions to contracting directed paths via direction-preserving homotopies and to the compatibility of path compositions [52–56]. In practice, path homologies provide a natural, context-independent description of serial activity pools and, together with simplicial homologies, help to clarify how neuronal computations may be organized. In the following sections, we explore this question by simulating the simplicial complexes of coactivity and the matching complexes of firing sequences produced by the hippocampus during spatial navigation, and then evaluating their spatial and ordinal structure and discussing the physiological implications of their homological classification.

## III. RESULTS

### A. Ordinal homology of assembly sequences during navigation

We simulated the rat’s movement in a small, low-dimensional enclosure covered by place fields, with positions sampled uniformly at random (Fig. 4A). From the resulting spiking patterns, we constructed the coactivity graph 𝒢 by connecting pairs of frequently cofiring cells and, for later comparison, evaluated the dynamics of the corresponding coactivity complex 𝒯 (Fig. 4B; see also [40, 42–44, 66]). At early stages of navigation, when only a few cell groups have had a chance to cofire, 𝒯 is small, fragmented, and contains numerous holes, cavities, tunnels, and other features, most of which lack physical counterparts. Under biologically realistic spiking parameters, these “spurious” structures tend to disappear as the pool of coactivities grows (Fig. 4C). Consequently, the complex 𝒯 becomes representable and takes on the topological shape of the surrounding environment [64]. In other words, as spiking accumulates, the pool of homologous simplicial chains consolidates, revealing the topological structure of the underlying space (Fig. 4B) [40, 42–49], which in turn is reflected in the animal’s spatial behavior [79–85].

**FIG. 4.**
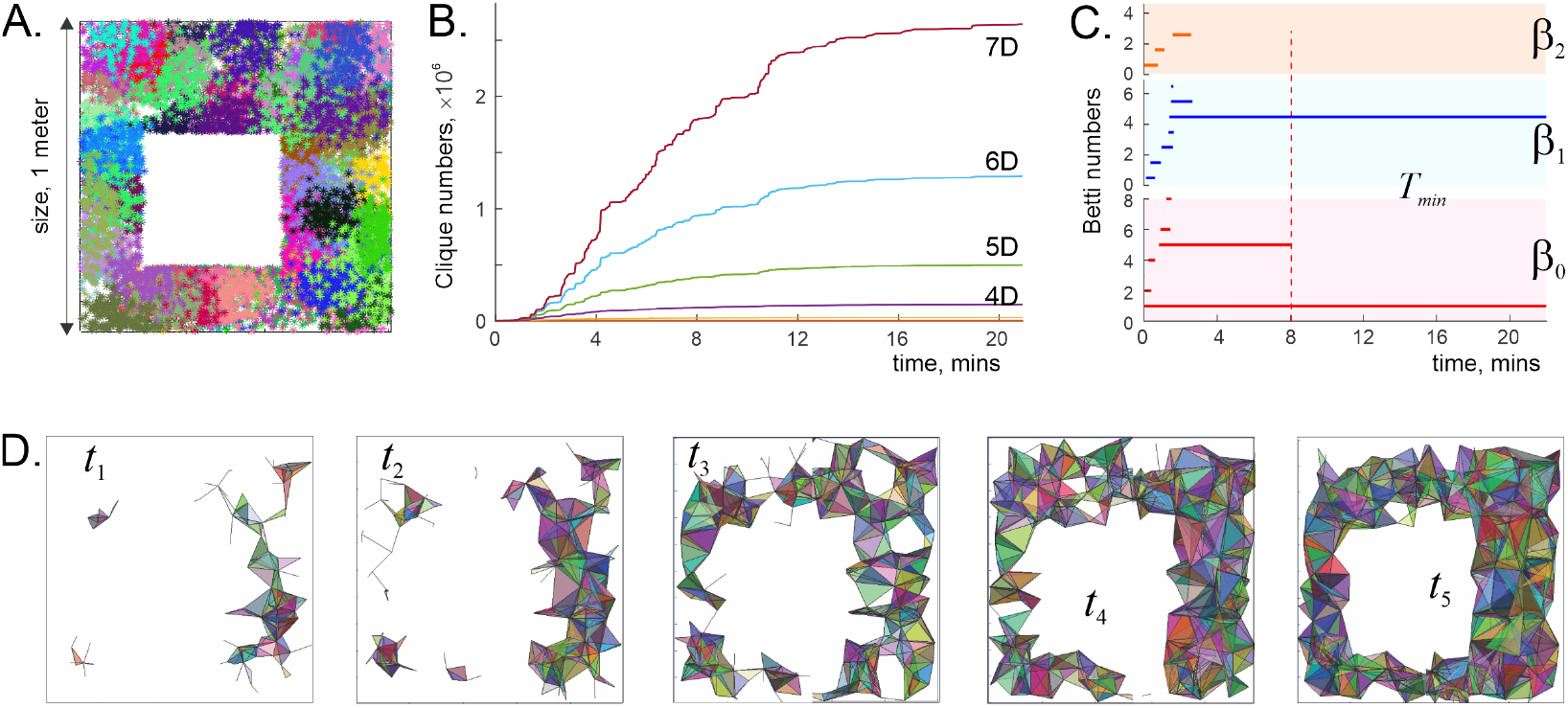
Simplicial dynamics. **A**. 2*D* arena with one hole, 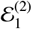, sized 1*m* × 1*m*, typical for electrophysiological experiments [88, 89], covered by 200 simulated place fields. The cells fired in the order their fields were traversed. Each dot of a particular color represents a spike fired by the corresponding cell at that location. Each cluster of colored dots represents a place field. **B**. As the animal navigates, the set of cells that have spiked grows, as does the pool of coactive cell combinations. Shown is the growing population of seven-dimensional (7*D*), six-dimensional (6*D*), five-dimensional (5*D*), *etc*., cliques of 𝒢, *i.e*., simplexes of 𝒯_𝒢_. **C**. The corresponding topological evolution of the coactivity complex: at first, 𝒯 is small and fragmented, but as the complex grows in size, its structure simplifies. Top, orange streaks mark the timelines of the noncontractible 2*D* cavities, middle, blue timelines pertain to 1*D* holes and the bottom, red timelines follow 0*D* loops—pieces of 𝒯. From early on, higher-dimensional Betti numbers, β_*D*≥2_, are suppressed, allowing representability of the complex [64]. The spurious holes close up by *T*_min_(𝒯) = 3.2 min, and complex fuses into one piece in about 8.0 mins, at which point 𝒯 becomes topologically equivalent to 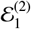. **D**. Pictorial dynamics of the coactivity complex, with 2*D* facets projected into the navigated environment, at five consecutive times (*t*_1_ ≈ 1 m, *t*_1_ ≈ 2 m, *t*_3_ ≈ 4 m, *t*_4_ ≈ 5 m, *t*_5_ ≈ 7 m).

On the other hand, the very same process can be interpreted as the formation of a path complex of simplicial trajectories. Indeed, the simplicial complex 𝒯 may be viewed as one or several “folded” simplicial paths: the assemblies that ignite at the animal’s current position produce active simplexes at the “tip” of an unfolding simplicial path Γ, while the simplexes left behind in its “tail” accumulate into 𝒯.

Furthermore, at each moment of its development, 𝒯(*t*) can be represented by the maximal simplex connectivity graph 𝔊_𝒯_ (*t*), whose vertices correspond to cell assemblies—the maximal simplices *σ*_*i*_ ∈ 𝒯(*t*)—and in which two vertices are connected by an edge *e*_*ij*_ if *σ*_*i*_ overlaps with *σ*_*j*_. The series of igniting cell assemblies then corresponds to sequences of edges in 𝔊_𝒯_. In the spirit of the persistence approach [86, 87], the path homologies of this evolving graph reveal, moment by moment, how the ordinal structure of igniting cell-assembly series unfolds over time. The corresponding homological dynamics may also modulate the animal’s behavior and cognitive performance, as one would theoretically expect.

For the neuronal ensemble with the place field map illustrated in Fig. 4A, the graph 𝔊_𝒯_ initially appears fragmented, as indicated by large values of the path-homological 0^th^-order Betti numbers, β_0_(𝔊_𝒯_) ~ 8. This is expected, as the disconnected subgraphs correspond to the connected components of 𝒯; the 0^th^ order topological loops appear and disappear simultaneously, indicating the consolidation of spatial and ordinal coactivity domains, *β*_0_(𝒯) = β_0_(𝔊_𝒯_).

On the other hand, first-order homologies, *H*_1_(𝒯) and 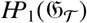, may differ significantly from each other. This is because 1*D* link-chains in 𝒯 “cycle” relative to their simplex boundaries (4) differently than the chains of 𝔊_𝒯_ edges “cycle” relative to the path boundaries (7). In particular, the high computational cost of evaluating path homologies (Fig.5B) limits the sizes and of the coactivity graphs that can be used in constructing path complexes, *i.e*., the sizes of spiking populations. Therefore, we restricted our analyses to the series of the most prominently firing cells—those that produce at least four spikes per burst (Fig.2E, [77, 78]).

The results show that the homological dynamics of the emerging path complexes, 𝒫_𝔊_(*t*), share several similarities with the dynamics of the coactivity complex 𝒯_𝒢_ (*t*): the population of distinct 1*D* cycles initially increases to high values, then decreases and stabilizes as distinct serial patterns become path-homologous. This happens over a path-learning period, *T*_min_(𝒫), that slightly exceeds the simplicial learning time, *T*_min_(𝒯). A similar trend is observed in the population of 2*D* loops, while higher-order Betti numbers, β_*k*>2_, tend to vanish. The latter indicates that homologically distinct chains composed of long (*k* > 3) elementary paths tend to consolidate. In plain words, longer sequences of assemblies ignited by traversing firing field maps tend to be structurally equivalent to one another, thus capturing less information, whereas collections of shorter sequences form richer patterns.

Importantly, this phenomenon is not generic: in our simulations, the path-homological Betti numbers of random graphs (constructed by randomly adding links between vertices) tend to increase with graph size [90], suggesting that the “homological equalizing” of longer cell-assembly sequences may be a specific property of map-representing graphs.

Adding directionalities to the edges of 𝔊_𝒯_, *i.e*., simulating preferred directions of activity propagation, does not qualitatively alter the previous scenario. The behavior of β_0_ remains qualitatively unchanged, and β_1_ shows exuberant dynamics at the early stages of navigation, which then subsides (longer timescales, with lower intermediate values).

Sequences of neuronal activity traced in most electrophysiological studies follow the connectivity of the original cognitive graph 𝒢, where vertices correspond to individual cells and links represent functional or synaptic connections between them. This graph is typically sparser than 𝔊_𝒯_, which relaxes the numerical restrictions on firing activity. We simulated the development of this graph during the rat’s running session discussed above (Fig. 4A), focusing on cells that produced at least three spikes per burst (Fig. 6A). The resulting path-homological dynamics exhibited familiar characteristics: initially, 𝒢 induces many spurious, heterologous sequences, which is followed by a period of consolidation and stabilization (Fig. 6B).

The fact that this period aligns closely with the cell assembly sequence learning timescale suggests that information encoded by both the neuronal and cell assembly sequences becomes accessible over similar timeframes. Additionally, the number of homologically distinct, closed neuronal firing sequences is comparable to the number of heterologous cell assembly path cycles and the number of simplicial loops in 𝒯 (Figs. 4, 5). If the connectivity graph is directed, reflecting, *e.g*., the synaptic connections between neurons, then the intermediate population of nonidentical short sequences is higher (β_1_ and β_2_ grow), but the overall path-homological dynamics remains similar.

**FIG. 5.**
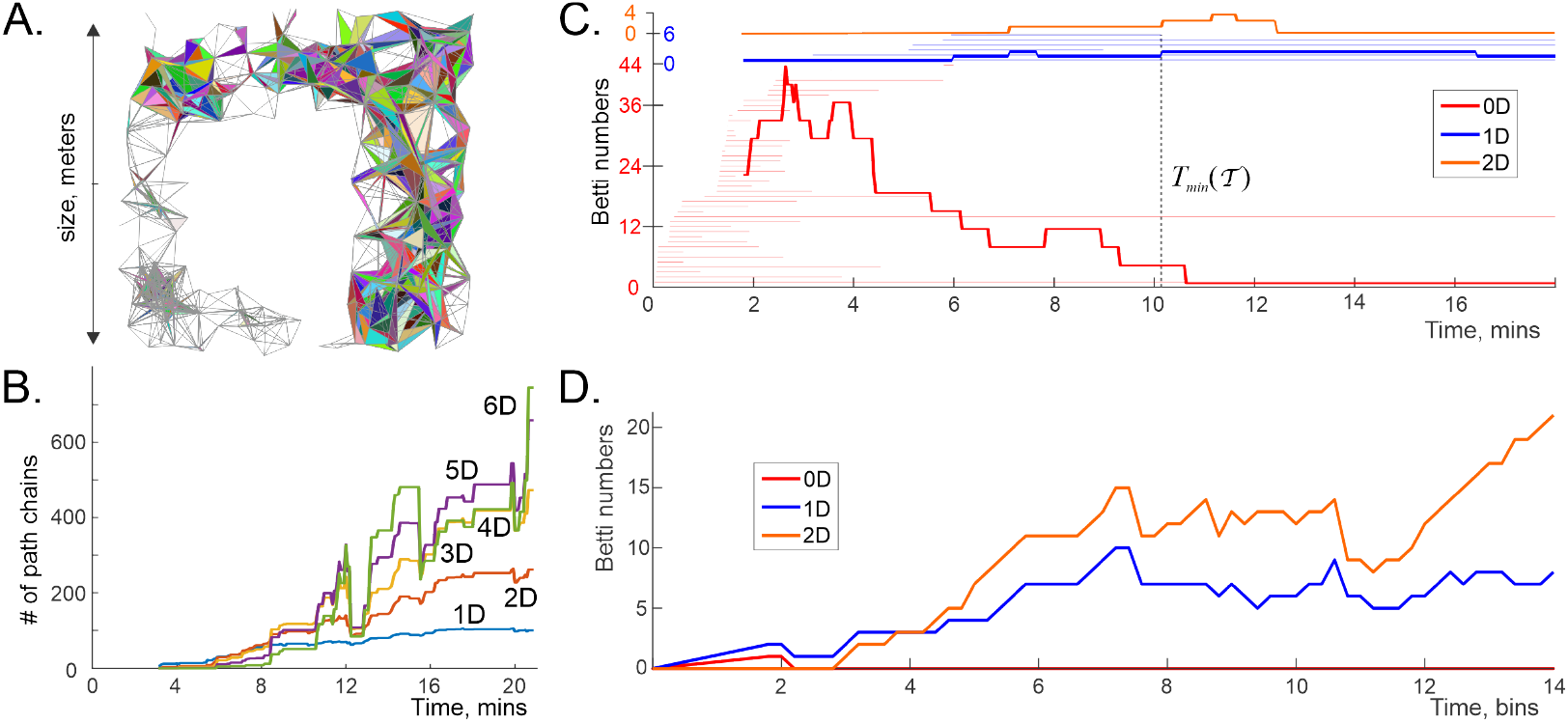
Ordinal structure of cell assembly sequences. **A**. Illustrative representation of the connectivity graph 𝔊_𝒯_ of the coactivity complex 𝒯_𝒢_ (gray bonds), defining incidences between maximal simplices at a given stage of its development. **B**. The population of 5-vertex, 4-vertex, 3-vertex, *etc*., path-chains in the path complex 𝒫_𝔊_, induced by the maximal simplex connectivity graph 𝔊_𝒯_, grows over time. **C**. The path-homological dynamics of 𝒫_𝔊_ exhibits similarities with the simplicial-homological dynamics of 𝒯_𝒢_: the number of distinct path-cycles initially increases and then decreases over time. The plot shows the numbers of 0*D* (blue, β_0_), 1*D* (red, β_1_), and 2*D* (orange, β_2_) pathcycles. Higher-order path-chains do not produce additional homology classes (β_*k*>2_ = 0). The horizontal lines in the background indicate the lifetimes of spurious topological loops in 𝒯_𝒢_, identified by simplicial persistent-homology analyses (see Fig. 4C, [40]). **D**. In a growing random graph (connected, β_0_ = 1), the number of path-homologically distinct path-cycles (path-homological Betti numbers) increases, but the number of heterologous chains also remains high (*D* ≳ 6, not shown).

In a 3*D* environment, connectivity graphs derived from place field maps yield an ensemble of homologically distinct 2*D* path loops, present in both vertex sequences (series of igniting neurons) and clique sequences (series of igniting cell assemblies), persisting throughout the simulated period. To obtain these results, we simulated movements through a 3*D* topological cylinder over the 2*D* environment shown on Fig. 4A, 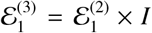, where *I* = [0, *L*] is a Euclidean interval (Fig. 6C). The size of the base and the height, *L*, were scaled to match the proportions of a bat’s cave described in [91]—about 3 meters across [43]. The resulting space has the same fundamental group, 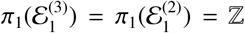, and the same topological complexity [92, 93], and therefore poses a comparable topological learning task as its 2*D* counterpart. The lengthening of the path-homologically nontrivial spiking series may hence be attributed to the increase of the underlying space’s dimensionality (Fig. 6D).

To investigate whether the maximal length of path-homologically distinct sequences,

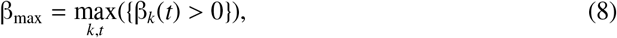

and its duration, |*T* |, where

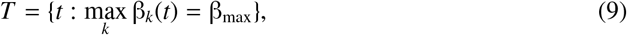

continue to increase with the dimensionality of the underlying space, *D* = dim(ℰ)—a dynamic also observed in the simplicial homologies of the coactivity complex [64]—we leveraged the fact that all components of our construction (trajectory, place fields, etc.) can be extended to any dimension without affecting the topological complexity or the homotopical and simplicial-homological structure of the space. Correspondingly, we constructed several coactivity graphs from the place field maps covering a 4*D* analogue of the 3*D* cave (Fig. 6C) and 2*D* navigational arena (Fig. 4A), 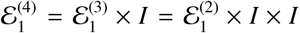, and observed that the *path-homological Leray index* (8) increased to β_max_ = 3. The emerging β_max_ ∝ *D* trend in map-induced path complexes (Fig. 6E), as well as the general suppression of higher-order Betti numbers requires further investigations. Biologically, this would imply that the homogenization of longer sequences and the richness of shorter sequences depend on the complexity of the context into which the network is submerged.

### B. Memory spaces

Episodic memory frameworks can be regarded as spaces in which specific memories, *m*_1_, *m*_2_,…, correspond to regions, *r*_1_, *r*_2_, … [94]. Neurophysiologically, this perspective is instrumental because it allows viewing many cognitive phenomena—such as spatial planning and exploration, transitive and relational inference, memory retrieval, and others—as particular cases of “mental navigation,” thereby facilitating physiological interpretations of data [94–100].

From a topological standpoint, this interpretation is also natural, as simplicial complexes define finitary spaces: individual simplexes emulate points, *x*_*i*_ (“memory nodes”), and their agglomerates represent broader memory scopes, *r*_*i*_. Essentially, this framework can be viewed as a physicalist extension of the cell assembly theory [22, 23], which associates cognitive episodes with the activity of cell assemblies. It postulates mappings from coactivity complexes, 𝒯, into a cognitive map or memory space,

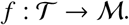

The totality of such mappings defines a *singular coactivity complex* associated with ℳ, whose homologies determine its topological structure [100].

As mentioned above, a remarkable property of hippocampal coactivity complexes is that they are *representable*, meaning there exists a domain in a low-dimensional Euclidean space, *X*, covered by a set of regular regions, *υ*_1_, *υ*_2_, …, *υ*_*N*_, whose nonempty overlaps correspond one-to-one with the coactivity simplexes. Specifically, 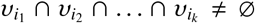 if and only if 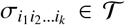, [64, 65]. As discussed *in [64]*, it is the dimensionality *of X* that limits the order of non-vanishing Betti numbers, *β*s. The experimental discovery of hippocampal representability—the existence of place fields marked a major advance in our understanding of learning, memory, and cognition. In particular, the match between Č ech homologies of the place field maps and the simplicial homologies of the coactivity complex supports interpreting the latter as a discrete topological map of the navigated space [101–103]. Similarly, the match between the simplicial homologies of place cell coactivity and the singular homologies of the corresponding discrete Alexandrov space suggests viewing the latter as a discretized representation of the environment embedded within memory space ℳ (Fig. 4A) [104–108].

The possibility of describing the ordinal structure of episodic memories by classifying sequences of regions distributed over ℳ,

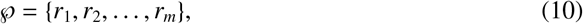

opens a complementary avenue for these analyses. Structurally, the “topological fabric” of a modeled memory space is defined by the pattern of immediate (minimal) neighborhoods, *U*_*i*_, of its points, which is described by the *topogenous matrix* [109]:

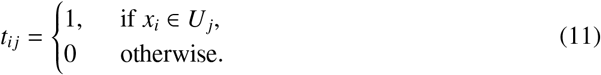

Here, we assume that the memory spaces are *T*_0_-separable, meaning that different points, *x*_*i*_ ≠ *x*_*j*_, have distinct minimal neighborhoods, *U*_*i*_ ≠ *U*_*j*_, and we set *t*_*ii*_ = 0 [104–108]. The matrix (11) can then be viewed as the connectivity matrix of a *topogenous graph* 𝒢, where vertices represent elementary memory nodes and edges define topological overlaps between them.^4^ The path homologies of the graph 𝒢 can thus describe the ordinal organization of memory sequences.

To test its properties, we constructed the graph 𝒢 for the Alexandrov spaces corresponding to the snapshots of the evolving coactivity complex 𝒯(*t*), focusing on the most prominently firing cells (at least six spikes per burst period), and traced their path homologies at different stages of the memory spaces’ development. Once again, the dynamics of β_*k*_(𝒢) largely mirror those of β_*k*_(𝔊) and β_*k*_(𝒢), suggesting that the emerging episodic memory chains encoded by place cell activity are structurally similar to cell assembly and neuronal chain order dynamics: disordered at first, then consolidating over the path learning period *T*_min_(𝒢).

It is worth noting that the pools of memory sequences could also be analyzed using singular-like homologies, in terms of chains induced by mapping 𝔊_𝒯_ into 𝒢, though we do not pursue this approach here [54].

### C. Topological consolidation

As discussed in the Introduction, the hippocampus generates coarse representations of the external world—a “subway map” of memorized experiences [111, 112]. Homological descriptions of this map are even coarser: the network’s structure and spiking dynamics may change significantly without affecting the path or simplicial homologies of the neuronal coactivity complex. For instance, alterations in the simplicial complex 𝒯, such as contractible protrusions, bulges, bends, or overall enlargements or shrinkages (Fig. 4C), remain indiscernible in homological analyses as long as no topological loops are created or eliminated. In other words, homological descriptions reflect only major, qualitative restructurings. Yet, physiological changes can be subtler, such as the consolidation of rapidly learned hippocampal memories into longer-term, coarser-grained cortical structures that preserve the spatiality and overall morphology of the original maps [113– 116, 118, 119, 130].

The exact physiological mechanisms underlying these processes are unclear, but functionally, they involve eliminating redundant or disused cell assemblies and consolidating inputs from active ones [120, 121]. Phenomenologically, such effects can be modeled by removing structurally redundant simplexes from the coactivity complex 𝒯, reducing the granularity of the corresponding finitary topological spaces while preserving their homologies [56]. In particular, the maximally coarsened memory space—the *core*, 𝒞(ℳ)—captures the original topological structure using the smallest number of points and neighborhoods (Fig. 7A) [104–108]. Interestingly, similar compact representations are also discussed in neuroscience as Morris’ “location schemas” [72, 122–124]. In [100], it was argued that such schemas can be constructed as the cores of memory spaces produced by place cells in a given environment, for specific physiological parameters of neuronal activity.

**FIG. 6.**
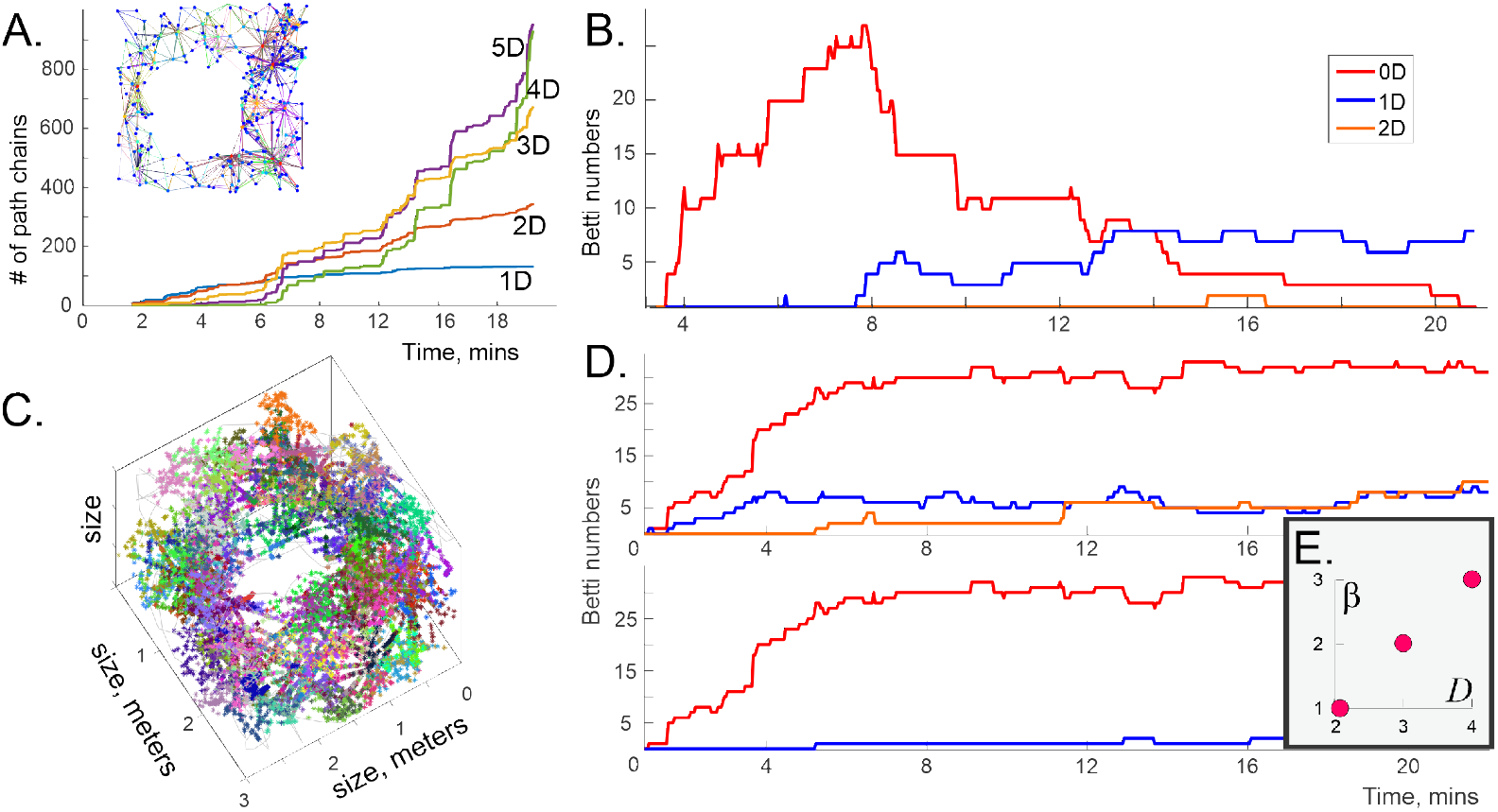
Ordinal structure of neuronal sequences. **A**. The population of 5*D*, 4*D*, 3*D*, … path-chains in the path complex induced by the growing cognitive graph 𝒢 (embedded panel) grows with time. **B**. Path-homological dynamics of the coactivity graph 𝒢 (neuronal sequences) is consistent with the dynamics of 𝒫_𝔊_ (cell assembly sequences): the number of distinct path cycles (Betti numbers β_*k*_) first grows and then decreases with time. Higher-order path cycles vanish. **C**. Place field map in a 3*D* environment with one hole, 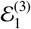 (simulated bat movements [43]). Points correspond to spikes fired by a specific cell at a particular location. **D**. 3*D* navigation produces more path-homologically distinct length-0 path cycles, which means that for the select size of place cell ensemble, speed value and the spiking parameters, the coactivity graphs (undirected, top and directed, bottom) have many disconnected components. Note also that the undirected coactivity graph generates a population of persistent, homologically distinct length-2 cycles, which appear only fleetingly in panel B. **E**. Lasting (|*T* | ≳ *T*_min_(𝒫)) path-homological Betti-order of the undirected cell assembly connectivity graph grows with the dimensionality of the navigated space.

**FIG. 7.**
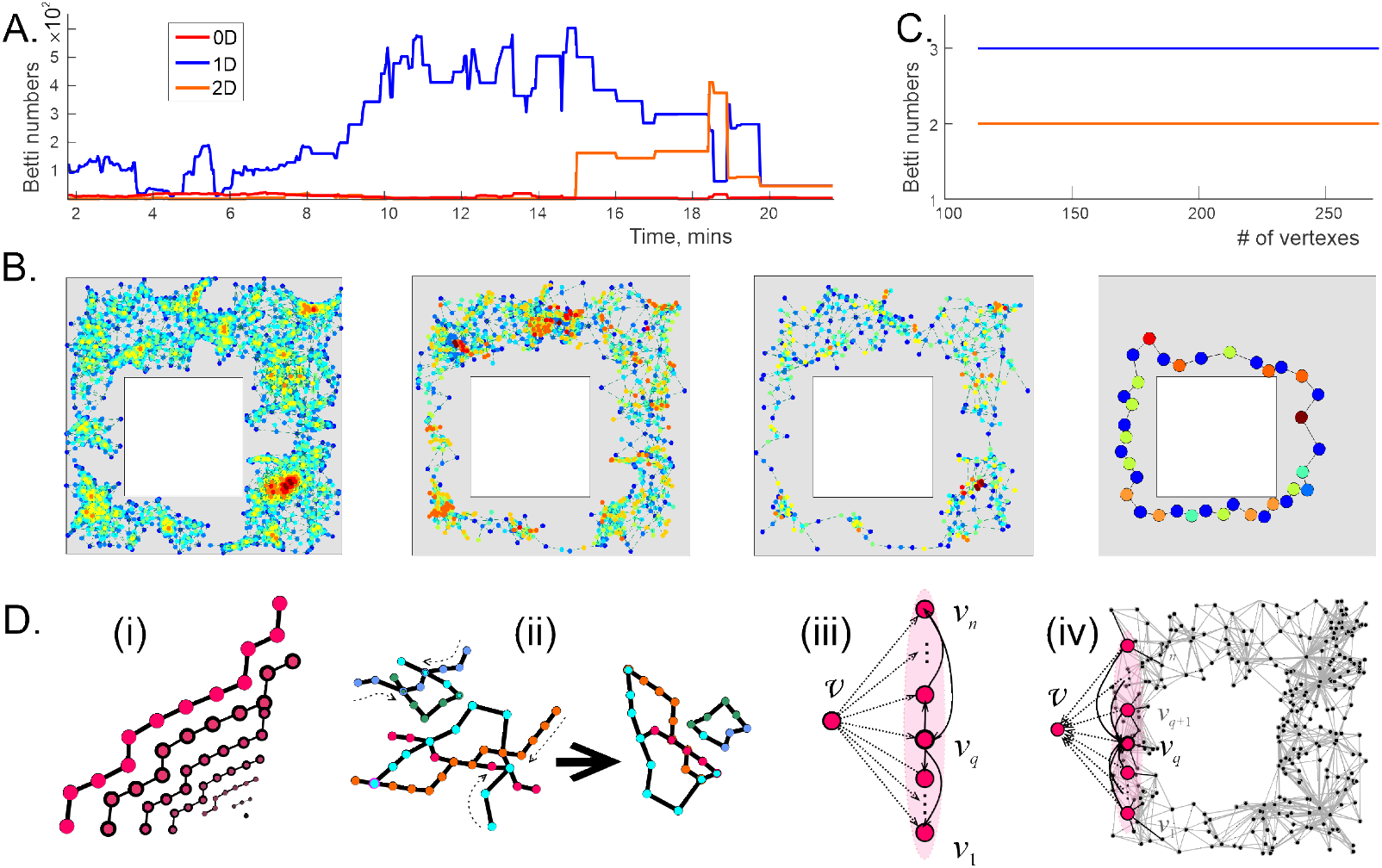
Memory template contraction. **A**. Path-homological Betti numbers for a series of topogenous graphs associated with the Alexandrov spaces induced at different stages of the coactivity complex’s development. The topological dynamics, as described by simplicial and singular homology, are identical to those shown in Fig. 4A. **B**. Elementary locations of the Alexandrov space built for the place field map shown on Fig. 4A, and the corresponding location maps for several of its retractions (maps folding a space onto a subspace; see Sec. VI). The rightmost panel shows the core, a maximally reduced schema of the memory space containing only 36 locations. Colors indicate the dimensionalities of the simplices giving rise to the points [45, 110]. **C**. Path-homological Betti numbers computed for the topogenous graphs of the retracting Alexandrov space remain unchanged, suggest that topological retraction also induces a homotopy equivalence of the corresponding path complexes. **D**. Illustrations of path homotopies. (i) A linear path can be contracted to a single node. (ii) “Tendrils” extending out of a generic path complex can be retracted. (iii) Star graph: if a single “apex” vertex v connects to all other vertices *v*_1_, *v*_2_, …, *v*_*n*_, either by outgoing (shown) or by incoming edges, then the entire graph can be contracted to v, so that all of its path homology groups vanish. (iv) If a vertex v connects to a set of vertices *v*_1_, *v*_2_, …, *v*_*k*_, and if one of these vertices, say *v*_*s*_, connects to the others exactly as v does (*i.e*., 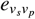 exists whenever 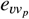 exists and has the same orientation, for *p* = 1, …, *k*), then v together with all its adjacent edges can be removed from the graph *G* without changing its path homology.

From the path-homological perspective, coarsening a memory space implies a reduction in the associated pool of memory sequences, which could potentially disrupt the overall structure of serial activity. However, direct computations reveal that the path homologies of the “consolidated” set of sequences remain unchanged—that is, a smaller population of heterologous sequences in reduced memory spaces preserve the structure of the original pool (Fig. 7B). In other words, patterns of consolidated activity retain both their ordinal and spatial organization. Biologically, this suggests that serial and spatial frameworks can be consolidated consistently.

Mathematically, the preservation of memory sequences’ homologies results from the fact that spatial reduction amounts to a homotopical transformation of the path complex. By this reasoning, one can use generic graph-homotopy retractions to model the consolidation of the sequential activity framework and the associated decimations of the neuronal population. These transformations, by preserving path homologies, yield a smaller repertoire of spiking activity that retains the original memory structure over smaller memory templates.

For instance, isolated sequences can be contracted to a single node, indicating that, at the level of abstraction used in this approach, solitary series of cognitive events may consolidate into a single episode (Fig.7A,B). Similarly, linear ‘tendrils’ extending from the graph can be retracted to its base, suggesting that sequences of extraneous cognitive episodes can be integrated into a compact core (Fig.7D). By this reasoning, sequential memory graph schemas [125] can in general be modeled as retracts of the corresponding coactivity graphs.

## IV. DISCUSSION

Representing spiking patterns by simplicial coactivity complexes serves several aims. First, it provides a straightforward qualitative model of the cell-assembly network—a generic schema that can incorporate numerous cellular-level characteristics and reveal their system-level effects [45, 47, 48, 125]. Second, a coactivity complex 𝒯 can be viewed as the Č ech complex associated with the place field map. In morphing environments, where this map deforms “elastically” [31–37], its Č ech complex remains intact, underscoring the topological nature of the encoded information and the stability of the network configuration that produces this map. Third, the simplicial semantics highlight a particular organization of network activity by identifying equivalence classes of igniting cell-assembly patterns and interpreting them as homologous chains of simplexes. Given the hippocampus’ role in establishing the topological aspect of spatial awareness, such equivalences may be innate to the hippocampo-cortical network [79–85].

On the other hand, recent studies portray the hippocampus as a universal generator of firing sequences, which act as functional motifs triggering cognitive episodes and behaviors [25, 126, 127]. Experimental investigations into sequential neuronal firing reveal familiar adaptability: the same firing order can adapt to different spatial and temporal scales, depending on specific tasks, contexts, or environments [27–30]. In physical terms, a particular configuration of firing sequences may also reflect a quasi-stable state of the underlying network, that can be schematically represented by a graph, dynamically manifested as a path complex [125]. The intrinsic arrangement of this complex’ components is captured by an alternative notion of equivalence between its components— path homology, that permits classifying physiologically distinct sequences—igniting assemblies, firing cells, evoked memory episodes, *etc*. This approach allows categorizing directed paths, tracing how they concatenate, identifying obstructions to contraction and characterizing the fragmentation of sequence pools according to the permissible transformations between sequences.

The results suggest, *e.g*., that hippocampal sequences generated by hundreds of cells and cell assemblies fall into only a few equivalence classes that collectively define the ordinal organization of hippocampal activity. Furthermore, it sustains the retractions of the coactivity complex, pointing to the possibility of simultaneous consolidation of ordinal and spatial maps [128]. Path-homological analyses also indicate that ordinal structures are predominantly supported by shorter sequences, much like spatial maps rely on the coactivity of smaller cell groups [45, 110], suggesting that biological information is generally conveyed by small computational units. Finally, the ordinal organization of neuronal firing emerges and stabilizes over timescales comparable to those of spatial map learning, exhibiting similar dynamics.

Certain parallels between simplicial and path-homological descriptions of hippocampal dynamics are natural: both offer complementary views of the information flow through the same network, share building blocks, and are functionally interconnected. For example, hippocampal maps shape spiking order (much as subway maps constrain the order in which stations are visited), and, conversely, expanding pools of interwoven sequences can form nexuses of consistently linked elements that yield connectivity maps (just as traversing enough stations reveals the metro plan).

It might therefore seem that the differences between these perspectives are merely semantic, *e.g*., if the learning process is viewed as spatial mapping, homologous spiking patterns would naturally be identified simplicially, whereas if the emphasis is on sequential experience, such as memorizing routes between locations, then tracing path homologies among ordered activities would be more appropriate. However, this is not naïvely the case: the principles that define homology in these two approaches differ qualitatively, which leads to fundamentally distinct calculations, with different computational costs. Ordinal structures do not, in general, reflect spatial affinities and vice versa, leading to different time scales over which these structures emerge. In other words, the spatial and ordinal cognitive frameworks are not interchangeable: the task of “mapping” a new environment is distinct from the task of memorizing routes or other sequential experiences. Thus, the ongoing “space vs. sequences” debate in the hippocampal literature has a substantive mathematical core [25, 38, 75, 76, 129].

At present, it remains unclear which homological framework best captures the essence of physiological computation; this must be determined experimentally. The prominence of near-simultaneous ignitions of cell groups [9–12] and of serial firing motifs [74, 132] suggests that both principles are implemented in the hippocampus and/or supporting networks. In either case, the proposed two-sided homological description provides a principled generalization of the hypothesized topological coding in the hippocampus [37, 38, 40, 80].

Many physical systems exhibit structures and dynamics that arise from homological and cohomological equivalences among their components [130, 131]. Similar principles may govern neuronal systems, where simplicial or path-homological equivalences could define large-scale, system-level organization of spiking patterns in ways that manifest at the cognitive level. Indeed, representatives of homology classes often correspond to behavioral variables. For example, persistent simplicial *H*_1_-cycles may map to physical barriers or, in abstract tasks, to incompatible feature conjunctions. In the language of Č ech homology, this corresponds to place or feature fields that overlap pairwise around obstacles or boundaries, yet do not form chains that traverse them. In general, simplicial homology captures global voids of coactivity that reflect how physiological, physical, or cognitive constraints shape information flow through the network. Correspondingly, changes in homological structure induced by topological reorganization of the environment alter this flow, producing measurable behavioral and cognitive effects [79–85].

It is therefore natural to hypothesize that path-homological characteristics may also couple to physiological parameters: the network may respond to *ordinal* reorganization of the environment or task—even when such changes are not topological in the direct sense. In other words, qualitative changes in the order of neuronal spiking should elicit consistent, statistically significant behavioral responses reflecting adaptation to the altered serial structure—a prediction that can be tested experimentally.

## V. ACKNOWLEDGMENTS

We are grateful to Prof. A. Grigor’yan for the software used to compute the path homologies. Additional MATLAB software by Dr. M. Yutin is available from SteveHuntsmanBAESystems, see [133].

The work was supported by NSF grant 1901338 and partly by NIH grants R01AG074226 and R01NS110806.

## VI. MATHEMATICAL SUPPLEMENT: BASIC NOTIONS AND GLOSSARY

### 1. Geometric simplexes and complexes

- *Geometric d-dimensional simplex* κ^*d*^ is the convex hull of its (*d* + 1) vertices (Fig. 1A).
- The *facet* opposite to a given vertex in a *d*-dimensional simplex κ^*d*^ is itself a (*d* − 1)-dimensional simplex κ^*d*−1^ and vice versa, κ^*d*^ can be produced by joining the points of κ^*d*−1^ to a new, (*d* + 1)^st^ vertex (Fig. 1B). The order of vertices defines the *orientation* of a simplex.
- *Boundary of a d-simplex, ∂* κ^*d*^, consists of its (*d* − 1)-dimensional faces (Fig. 1B).

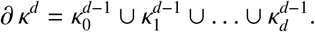
- A simplex is *maximal*, if it is not a subsimplex of any other simplex.
- *Geometric simplicial complex K* is a combination of properly assembled geometric simplexes that fit each other vertex-to-vertex, side-to-side, with no overruns or other mismatches (Fig. 1C). Formally, it is required that a non-empty intersection of any two simplexes *κ*_1_, *κ*_2_ in *K* is another simplex from *K*.
- The collection of all simplexes of dimensionality *d* and less forms the *d-skeleton* of a simplicial complex *K, sk*_*d*_(*K*).
- Any undirected graph is a 1*D* simplicial complex.

### 2. Abstract simplexes and complexes

- Since all that matters for quantifying topology of simplicial complexes is how their simplexes match, it is natural to ignore their “filling” altogether and pass on to the notion of *abstract simplexes*—ordered lists of simplexes’ vertices (1), and their collections— *abstract simplicial complexes*, Σ. An abstract simplex may be intuitively apprehended as the set of vertices of a geometric simplex. The “face-matching” of the geometric simplexes reduces to the requirement that if an abstract simplex belongs to a simplicial complex, then so do all its faces [58].

### 3. Graphs

- *Directed graph*, or *digraph, G* consists of vertices, *V* = { *v*_1_, *v*_2_, …, *v*_*n*_ }, and directed edges between certain pairs of vertices. Formally, *G* is described by its connectivity matrix

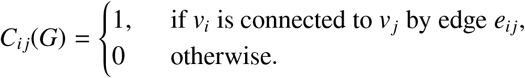
- If the matrix *C*_*ij*_ is symmetric, the graph is viewed as undirected, *i.e*., *e*_*ji*_ always exists if *e*_*ij*_ does.
- *Digraph map f* : *G* → *H* sends vertices *v*_*i*_ of the digraph *G* into the vertices *w*_*p*_ of the digraph *H*, so that each *G*-edge either maps on a *H*-edge, or squeezes into a *H*-vertex. Incidentally, digraphs and maps between them form a category.
- *G* is *transitive*, if whenever edges *e*_*ij*_ and *e*_*jk*_ exist, then *e*_*ik*_ is also an edge.
- *Line digraph, I*_*n*_, consists of *n* ordered directed links between consecutive pairs of vertices, *e.g*., *I*_*n*_ = {*v*_0_ → *v*_1_ ← *v*_2_ → … ← *v*_*n*−1_ ← *v*_*n*_}. If the last vertex, *v*_*n*_, connects to the first, *v*_1_, then digraph is *cyclic, e.g*., *S*_*n*_ = { *v*_*n*_ →*v*_0_ ← *v*_1_→… ← *v*_*n*−1_ →*v*_*n*_ }.
- Line digraphs of length one are the two *directed dyads*: *I*^+^ = (𝓋 𝓋^’^) and *I*^−^ = (𝓋 𝓋^’^).
- An *acyclic digraph* has no directed cycles, *i.e*., no sequences *e*_*ij*_, *e*_*jl*_, …, *e*_*qi*_. Such digraphs allow topological ordering of vertexes (consistent with edge directions), and have partially ordered directed paths.
- *Cartesian product* of a digraph *G* with a vertex set *V* and a digraph *H* with its vertex set *W*, is a digraph *G* × *H* whose vertices have a *G*- and a *H*-component, (*v*_*i*_, *w*_*p*_). Vertices (*v*_*i*_, *w*_*p*_) and (*v*_*j*_, *w*_*q*_) are connected by an edge if either *v*_*i*_ connects to *v*_*j*_ in *G*, with the *H*-component fixed, *w*_*p*_ = *w*_*q*_, or vice versa, *w*_*p*_ connects to *w*_*q*_ in *H*, with fixed *v*_*i*_ = *v*_*j*_.
- *Cylinder* over a digraph *G* is the Cartesian product, *G* × *I*_*n*_.
- *Clique* of order *d, ς*^(*d*)^ is a fully interconnected subgraph with (*d* + 1) vertices 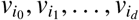.
- *Clique complex* associated with a graph *G*, Σ_*G*_, is the full collection of *G*-cliques [134].
- A finite simplicial complex Σ can be represented by the following constructions:
  i. vertices of the *simplex connectivity graph, G*_Σ_, *v*_*σ*_, represent simplexes of Σ, connected by an undirected edge *e*_*σσ*_′ if *σ* and *σ*′ intersect, *σ* ∩ *σ*^’^ ≠ ∅ [134].
  ii. vertices *v*_*σ*_ and *v*_*σ*_′ of the *maximal simplex connectivity graph*, 𝔊_Σ_, are connected by an undirected edge *e*_*σσ*_′ if the maximal simplexes *σ, σ*^’^ ∈ Σ intersect, *σ*^’^ ∩ *σ* ≠ ∅.
  iii. vertices *v*_*σ*_ and *v*_*σ*_′ of the *simplex inclusion digraph*, G_Σ_, are connected by a directed edge *e*_*σσ*_′ if *σ*^’^ is a facet of *σ, i.e*., if *σ* ⊊ *σ*^’^.
  iv. If Σ is a clique complex, then the *barycentric refinement* of its 1*D* skeleton, *sk*_1_(Σ) = *G*, is the graph whose vertices correspond to cliques of *ς* of Σ, and the edge *e*_*ςς*_′ between *ς* and *ς*^’^ exists if *ς* <; *ς*^’^. Thus, barycentric refinement of *G* is the connectivity graph of its clique complex Σ_*G*_.
  v. vertices *v*_*σ*_ and *v*_*σ*_′ of the *face connectivity digraph*, G_Σ_, connect by *e*_*σσ*_′ if *σ* is a maximal facet of *σ*^’^, dim(*σ*) = dim(*σ*^’^) + 1.
  vi. vertices *v*_*x*_ and *v*_*x*_′ of the *topogenous graph* 𝒢_Σ_ represent points in a topological space *X*, and the edges *e*_*xx*_′ represent their adjacency (one point lies in the other’s minimal open neighborhood).

### 4. A few bullet points of the simplicial homology theory —for more details see [58–60]

- *Chains* of dimensionality *k* are formal linear combinations of oriented *k*-dimensional simplexes, in which 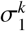 is counted *m*_1_ times, 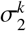 is counted *m*_2_ times, etc.,

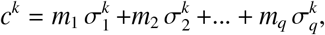

with the coefficients from a field 𝕂. If the coefficients come from the Boolean field ℤ_2_, then the chain is simply a list of “present” (*m*_*i*_ = 1) and “absent” (*m*_*i*_ = 0) simplexes. The chains can be added or subtracted from one another, as well as multiplied and divided by coefficients, thus producing linear spaces, *C*_*k*_(Σ). Intuitively, each chain space is an algebraization of its *k*-dimensional part, *e.g*., the dimensionality of *C*_*k*_ is given by the number of the *k*-simplexes in Σ,

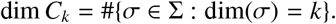
- *Boundary of a k-simplex*, algebro-topologically, is a (*k* − 1)-dimensional *chain of faces*, as defined by the formula (4). For example, the boundary of an interval *σ*_12_ oriented from vertex *σ*_1_ to vertex *σ*_2_ is a formal alternated sum *∂ σ*_01_ = *σ*_1_ − *σ*_0_. The boundary of a filled 2*D* triangle is an alternating sum of its three sides, *∂ σ*_012_ = *σ*_12_ − *σ*_02_ + *σ*_01_, etc.
- *Boundary of a chain* is a combination of the contributing simplexes’ algebraic boundaries,

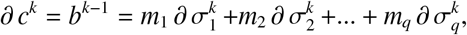

which is hence an element of the *C*_*k*−1_(Σ)-space. The full collection of boundary chains *b*^*k*−1^ forms a subspace of *C*_*k*−1_(Σ), denoted by *B*_*k*−1_(Σ).
- *Nilpotence*. By direct verification, *∂*^2^ *c*^*k*^ = 0, *i.e*., boundary of a boundary always vanishes.
- *Cycles*. Not all chains with vanishing boundary, *∂ c*^*k*^ = 0, are themselves boundaries. In general, such chains are combinations of simplex chains that “loop back onto themselves” within the complex. Such chains are referred to as *cycles*, and denoted by *z*. Linear combinations of cycles of dimensionality *k* also form a vector space, denoted by *Z*_*k*_(Σ), which is smaller than *C*_*k*_(Σ), but generally larger than *B*_*k*_(Σ) [58–60].
- *Chain complex* is a system of spaces *C*_*k*_, in which higher-order spaces are mapped into lower-order ones,

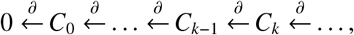

as the boundary of a *k*-dimensional chain is a (*k* − 1)-dimensional chain. In other words, at each step the boundary operator *∂* maps *C*_*k*_ into *B*_*k*−1_ that lays within *C*_*k*−1_.
- *Homologous chains*. Adding or removing a *k*-dimensional boundary amounts to deforming a cycle *z* by “snapping” it over a series of (*k* + 1)-dimensional simplexes and producing an equivalent, or *homologous* cycle *z* ± *∂ c* = *z*^’^ (Fig. 1D). Cycles that cannot be matched by adding or removing full boundaries are nonequivalent. Identifying equivalence classes hence amounts to factoring out the “boundary parts,” *i.e*., taking the quotient of the cycle space by the boundary space, *H*_*k*_(Σ) = *Z*_*k*_(Σ)/*B*_*k*_(Σ). The representatives of the resulting space of *k*^th^ *simplicial homology groups, H*_*k*_(Σ), represent *k*-dimensional loops, counted up to a continuous deformation.
- *Betti numbers* are the dimensionalities of homology spaces, *b*_*k*_(Σ) = dim *H*_*k*_(Σ); these numbers count the topologically distinct *k*-loops in the complex Σ. For example, deforming a 0*D* chain (a combination of 0*D* points) amounts to “sliding” those points inside of Σ; the dimensionality of the corresponding homology space (0^th^ Betti number), *β*_0_(Σ), is equal to the number of “sliding domains” in Σ, *i.e*., the number of its connected components. If a simplicial complex comes in one piece, its *β*_0_(Σ) = 1; *β*_1_(Σ) equals to the number of holes in Σ. An example of a 2*D* noncontractible loop captured by *β*_2_ is a “hollow” tetrahedron. Being topologically a 2-dimensional sphere, all 1-dimensional loops on it are contractible, and hence its Betti numbers are *β*_0_ = 1, *β*_1_ = 0, *β*_2_ = 1, *β*_*n*>2_ = 0.

### 5. Path homology theory is outlined below following the publications [52–56]

- Let *V* ={ *v*_1_, …, *v*_*n*_ } a finite set. An *elementary path* of length *k* is an arbitrary sequence of elements, 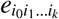. These paths are basic units of the theory, analogous to simplexes of simplicial topology.
- *Path complex* 𝒫 is the collection of the elementary paths. Lengths of paths define their order, *i.e*., correspond to the dimensionality of simplexes. We consider only finite path complexes that include paths up to some maximal length, *n*. If *P*_*l*_ is the set of all elementary paths of length *l*, then the collection, 𝒫_*k*_ = *P*_0_ ∪ *P*_1_ ∪ *P*_2_ ∪… ∪ *P*_*k*_, is the *k*-skeleton of 𝒫 —the analogue of the *k*-skeleton *sk*_*k*_(Σ) of a simplicial complex.
- The paths forming a complex 𝒫 must allow “plucking” of the endpoints, *i.e*., if 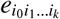 is in 𝒫, then 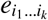 and 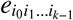 must also belong to 𝒫 (Fig. 2B).

### 6. Algebraization of path chain complexes

- *Path chains* of order *k* are formal linear combinations of elementary paths of length *k*, with the coefficients in a field 𝕂,

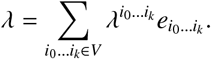

The collection of *all* path *k*-chains forms a linear space Λ_*k*_, *e.g*., Λ_0_ are linear combinations of vertices *e*_*i*_, Λ_1_, are the combinations of all pairs of vertices, *e*_*ij*_, etc.
- *Boundary operator* (7) breaks the elementary paths into segments that precede the *q*^th^ step and the segments that follow it, *e.g*., *∂ e*_*i*_ = *e, ∂ e*_*ij*_ = *e*_*j*_ − *e*_*i*_, *∂ e*_*ijk*_ = *e*_*jk*_ − *e*_*ik*_ + *e*_*ij*_, etc. By linearity, the boundary of a generic path chain is

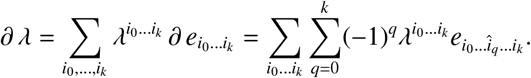
- By direct verification, *∂*^2^ *λ* = 0 for any *λ*, which implies that the boundary operator (7) induces a chain complex Λ_∗_(𝒫),

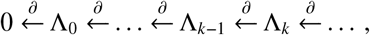

in which the path-chains nullified by *∂* are the path-cycles, *∂ ζ* = 0 and the chains of the form *β* = *∂ λ* are the boundaries. As with the simplicial complexes, both types of chains form their respective linear spaces, *Z*_*k*_(𝒫) and *B*_*k*_(𝒫). The quotient space of the path-cycles over the path-boundaries yield the *path homology space*, 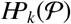, analogous to the simplicial homologies of simplicial complexes *H*_*k*_(Σ).
- If a path complex is formed by a collection of certain selected, or *allowed* elementary paths, then their linear combinations,

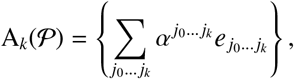

form a subspace of Λ_*k*_. Examples:

i. In a graph-representable complex, allowed paths are the ones that run along the graph’s edges, other sequence are excluded.
ii. The allowed paths on the (maximal) simplex connectivity graph are the sequences of overlapping (maximal) simplexes.
iii. From a biological perspective, if *V* is the full set of cells or assemblies in a given network, the allowed vertices, *P*_0_, may be the ones that exhibit activity in a given environment. The allowed links, *P*_1_, may be the ones that represent synaptic connections, rather than all pairs of coactive cells, *P*_2_, etc., may be synaptically interconnected triples of neurons, etc.

- In order for the restriction of the boundary operator (7) to *A*_*k*_, to be contained in *A*_*k*−1_, the boundaries of the allowed paths should also be allowed,

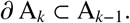
- However, in contrast with the simplicial case, where each term in the right hand side of (4) is a part of the complex Σ by design, terms appearing in the right hand sides of (7) may structurally fall out of 𝒫 (Fig. 2C). Yet, there exists a subclass of the allowed path chains— the *operational path-chains* that have allowed boundaries,

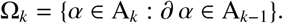

A simple, but fundamental property of such paths is that the boundary operator (7) acts on them without fallacies. Indeed, if *α* ∈ Ω_*k*_ then *∂ α* ∈ A_*k*−1_, and *∂*^2^ *α* = 0 ∈ A_*k*−2_, which means that *∂ α* ∈ Ω_*k*−1_. Thus, Ω-paths form their own chain complex, Ω_∗_,

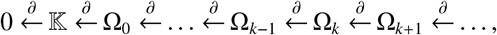

whose homology groups (the *reduced path homology groups* of 𝒫) define the structure of the allowed paths.
- An alternative way to view the action of the boundary operator (7) on *A*_*n*_ is to use the explicit projection,

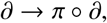

where *π* : Λ_*n*_ → *A*_*n*_ kills forbidden paths.
- A digraph map *f* : *G* → *H* induces a homomorphism of operational chain complexes,

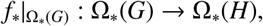

and of path homology groups,

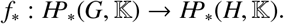

Thus, path homologies can be used to classify digraphs relative to such maps.

### 7. Simplicial vs. path homologies

- A path complex 𝒫 is:
  i. *perfect*, if it contains all subpaths of its elementary paths. In such complexes, the boundary (7) is structurally identical to (4).
  ii. *monotone*, if its vertices can be numbered such that the numbering increases along each elementary path in 𝒫. In a digraph, this implies that all edges comply with the ordering. For example, if a path complex contains two elementary paths with conflicting vertex order, *e.g*., 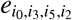 and 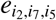 have a conflicting order of *i*_2_ and *i*_5_, then this complex clearly can not be monotone. Conversely, any finite complex without such conflicts is monotone.
  iii. perfect and representable by a digraph if and only if the latter is *transitive*.

- 𝒫 is the path complex of a simplicial complex if and only if it is perfect and monotone. This combination is rare in applications, *e.g*., the neuronal coactivity graph or the synaptic connectivity graph are typically not transitive. To build a positive example, consider elementary paths that pass through vertices enumerated in increasing order. The subsequences of such paths will also be ordered, *i.e*., the complex 𝒫_*G*_ will be perfect and representable by a transitive graph *G*(Σ), and the homologies 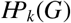 and *H*_*k*_(Σ), *k* ≥ 0, will be isomorphic [52, 53].
- in simplicial A “hole” means: there should be a big group that meets all together, but one is missing, leaving a gap in the friendship web.
- **Path homology** builds shapes out of *legal trips* you can actually drive in order (one-way roads). A “hole” means: there is a directed loop or a non-commuting detour you cannot shrink by allowed (one-way-respecting) moves.

### 8. Regularization of path complexes

- *Regular elementary paths* are the ones that contain no consecutively repeating indexes: in 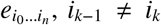 for all *k*. The chain spaces Λ_*k*_ can be decomposed into the regular and the irregular components, Λ_*k*_ = *R*_*k*_ ⨁ *I*_*k*_, which, however, do not produce separate path complexes, because *R*_*k*_ is not, by itself, invariant with respect to the boundary operator (7). For example, the boundary of a regular path *e*_*iji*_,

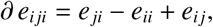

includes an irregular segment *e*_*ii*_. It turns out however, that one can build a homological classification of the regular paths simply by discounting “irregularities,” *i.e*., by viewing two path chains as equivalent, if they differ by an irregular path. In other words, one can use

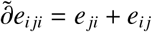

instead of the “naïve” *∂ e*_*iji*_ above. Each resulting equivalence class contains a unique regular path; that is, every path admits a unique regular representative, which we will henceforth call its *regularized* form. This procedure also induces a regularized boundary operator 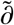, which allows defining the *regular chain complex*

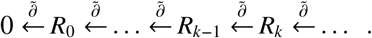

whose homologies describe regular paths [52].
- The set of allowed regular *k*-paths is a subset of the total set of regular *k*-paths. The boundary operator 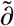 acting on such paths defines regular operational chains,

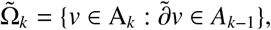

which are characterized by their regular homologies.
- A path complex 𝒫 is *strictly regular* if it is regular and does not breed irregularity, *i.e*., contains no paths of the form *e*_…*iji*_ For example, a path complex of a simplicial complex is strictly regular because its paths’ indices are strictly increasing. The path complex of a digraph is strictly regular iff there are no loops (no *e*_*ii*_ edges and no simultaneously present *e*_*ij*_ and *e*_*ji*_ edges). The corresponding homologies are the only ones used in our analyses.

### 9. Homotopies

- Two digraph maps *f, g* : *G* → *H* are *homotopic, f* ≃ *g*, if there is a line digraph *I*_*n*_ and a map *F* : *G* × *I*_*n*_ → *H* such that

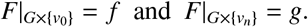

In particular, *f* and *g* are one-step homotopic (*n* = 1), *f* ≖ *g*, if *f* (*v*_*i*_) and *g*(*v*_*i*_) are either connected by an edge or coincide for all *v*_*i*_.
- Any homotopy, *f* ≃ *g*, amounts to a finite sequence of one-step homotopies, *f* = *f*_0_ ~_1_ *f*_1_ ~_1_ · · · ~_1_ *f*_*n*_ = *g*, where *f*_0_ = *id* is the identity map, and *f*_*k*_ ≖ *f*_*k*+1_.
- Two digraphs *G* and *H* are *homotopy equivalent, H* ≃ *G*, if there exist digraph maps *f* : *G* → *H* and *g* : *H* → *G*, such that

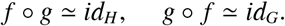

The maps *f* and *g* are then *homotopy inverses* of each other.
- Homotopy equivalent digraphs have isomorphic path homologies. Homotopic maps induce *identical* homomorphisms of path homologies.
  i. A cylinder over a digraph *G* is homotopic to its base, *G* × *I*_*n*_ ≃ *G*, for any *I*_*n*≥0_.
  ii. Natural inclusion of *G* into a cylinder *G* × *I*_*n*_ over it, *i*_*k*_ : *v* →(*v, ι*_*k*_), and the projection from a cylinder *G* × *I*_*n*_ to its base, *p* : (*v, ι*_*k*_) → *v*, induce isomorphism of path homologies.

- *Retraction* of a digraph *G* onto a subgraph *H* is a map *f* : *G* → *H* that fixes all the vertices of *H*. A *deformation retraction* is the one that can be homotopically deformed to the identity map. This implies that *G* and *H* are homotopically equivalent. Any deformation retraction is a composition of one-step retractions. A one-step retraction is a digraph map that fixes some vertices and moves all others either one step along the edges, or one step against the edges simultaneously.
- A digraph *G* is *contractible* if it is homotopic to a single vertex. All the homology groups of a contractible graph are trivial, except for 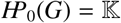 [135].
  i. A connected tree can be contracted to its root vertex by “trimming” the outer vertices. The process of removing the terminal vertices at each branch with its adjacent edge is a deformation retraction.
  ii. A *star-like* digraph has a vertex 𝓋 that connects to all other vertices by linear digraphs. Such a graph is contractible to 𝓋, *e.g*., a facet *σ*^(2)^ of a tetrahedron *σ*^(3)^ can be contracted to its opposite vertex.
  iii. A *k*-dimensional cube 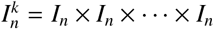 (direct product of *k* ≥ 1 line digraphs, *I*_*n*_), can be retracted to each of its facets, 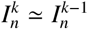, and ultimately contracted.
  iv. Cycling triangle and a cycling square are contractible. Longer cycles, *S*_*n*>4_, loose contractibility: *H*_1_(*S*_*n*>4_, 𝕂) ≅ 𝕂. Furthermore, two unequal cycles, *S*_*n*>4_ and *S*_*m*>4_, *n* ≠ *m*, are homotopy inequivalent.

- If *r* : *G* → *H* is a deformation retraction, then *G* and *H* are homotopy equivalent and the map *r* is homotopy inverse of *i* and vice versa.
- If a vertex 𝓋 connects to a group of other vertices, *v*_1_, …, *v*_*n*_, among which there is a special vertex, *v*_*p*_, that connects to all the remaining *v*s whenever 𝓋 does (𝓋 →*v*_*i*_ ⇒ *v*_*p*_ →*v*_*i*_), then removing 𝓋 along with all of its adjacent edges is a homology-preserving deformation retraction. This procedure may be viewed as removing this special kind of an “embedded star” from *G*. The arrows in this construction can be simultaneously reversed (𝓋 ← *v*_*i*_ ⇒ *v*_*p*_ ← *v*_*i*_).

#### 11. The topological complexity of a path-connected topological space

*X*, TC(*X*), is defined as the smallest number of open sets, *U*_1_, *U*_2_, … *U*_*k*_, required to cover *X* × *X*, such that each set *U*_*i*_ allows for continuous motion planning from a starting to an ending point within its domain [92]. The value TC(*X*) is a homotopy invariant [93], and for the spaces discussed in Sec. IIIA, 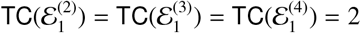.

## Footnotes

1 Throughout the text, semantic highlights and terminological definitions are given in *italics*. Key mathematical notions are outlined in Sec. VI

2 “…much like jazz musicians” [57].

3 In this study, we use the binary field, K = Z_2_ = {0, 1}, which is used without explicit reference.

4 The graph 𝒢 also describes connectivity between all simplexes in the coactivity complex.

